# Heme biosynthesis regulates BCAA catabolism and thermogenesis in brown adipose tissue

**DOI:** 10.1101/2023.11.28.568893

**Authors:** Dylan J. Duerre, Julia K. Hansen, Steven John, Annie Jen, Noah Carrillo, Hoang Bui, Yutong Bao, Matias Fabregat, Katherine Overmeyer, Evgenia Shishkova, Mark P. Keller, Richard A. Anderson, Vincent L. Cryns, Alan D. Attie, Joshua J. Coon, Jing Fan, Andrea Galmozzi

**Affiliations:** Department of Medicine, University of Wisconsin-Madison, School of Medicine and Public Health, Madison, WI; Cellular and Molecular Biology Graduate Program, University of Wisconsin-Madison, Madison, WI; Morgridge Institute for Research, Madison, WI; Integrated Program in Biochemistry, University of Wisconsin-Madison, Madison, WI; Department of Biomolecular Chemistry, University of Wisconsin-Madison, School of Medicine and Public Health, Madison, WI; Molecular and Environmental Toxicology Graduate Program, University of Wisconsin-Madison, Madison, WI; Metabolism and Nutrition Graduate Program, University of Wisconsin-Madison, Madison, WI; National Center for Quantitative Biology of Complex Systems, Madison, WI; Department of Biochemistry, University of Wisconsin-Madison, Madison, WI; Department of Chemistry, University of Wisconsin-Madison, Madison, WI; Department of Nutritional Sciences, University of Wisconsin-Madison, Madison, WI; University of Wisconsin Carbone Cancer Center, University of Wisconsin-Madison, School of Medicine and Public Health, Madison, WI

## Abstract

With age, people tend to accumulate body fat and reduce energy expenditure^1^. Brown (BAT) and beige adipose tissue dissipate heat and increase energy expenditure via the activity of the uncoupling protein UCP1 and other thermogenic futile cycles^2,3^. The activity of brown and beige depots inversely correlates with BMI and age^4–11^, suggesting that promoting thermogenesis may be an effective approach for combating age-related metabolic disease^12–15^. Heme is an enzyme cofactor and signaling molecule that we recently showed to regulate BAT function^16^. Here, we show that heme biosynthesis is the primary contributor to intracellular heme levels in brown adipocytes. Inhibition of heme biosynthesis leads to mitochondrial dysfunction and reduction in UCP1. Although supplementing heme can restore mitochondrial function in heme-synthesis-deficient cells, the downregulation of UCP1 persists due to the accumulation of the heme precursors, particularly propionyl-CoA, which is a product of branched-chain amino acids (BCAA) catabolism. Cold exposure promotes BCAA uptake in BAT, and defects in BCAA catabolism in this tissue hinder thermogenesis^17^. However, BCAAs’ contribution to the TCA cycle in BAT and WAT never exceeds 2% of total TCA flux^18^. Our work offers a way to integrate current literature by describing heme biosynthesis as an important metabolic sink for BCAAs.

## MAIN TEXT

How heme regulates adipocyte function and overall glucose homeostasis is poorly understood. Subjects with inborn errors of heme biosynthesis have a higher risk of developing glucose intolerance and insulin resistance^19^. Heme synthesis in mammals consists of an 8-step pathway^20^ that starts with the condensation of glycine and succinyl-CoA, catalyzed by the enzyme Aminolaevulinic Acid Synthase (ALAS) (**Extended Data Figure 1a**). Two distinct genes for the rate-limiting enzyme ALAS exist: one for erythroid cells (i.e., *ALAS2*) where heme is primarily synthesized for the production of hemoglobin, and one for all other cell types, *ALAS1*^21^. A testament to the importance of this metabolite besides its role in hemoglobin is that in mice, whole-body deletion of *Alas1*, the non-erythroid isoform, is embryonically lethal^22^. Conversely, whole-body *Alas1* heterozygous mice are viable but manifest accelerated age-associated impairments in glucose tolerance and mitochondrial function in skeletal muscle^23^. Furthermore, heme levels and the expression of *ALAS1 and ALAD*, the first and second enzymes of heme biosynthesis, are significantly reduced in the adipose of obese and diabetic subjects^24^, suggesting a link between heme and glucose homeostasis. In mice, *Alas1* expression varies widely among tissues, with the highest expression in the liver and adrenal gland. Notably, brown adipose tissue exhibits *Alas1* expression comparable to that of the heart and skeletal muscle and is significantly higher than inguinal and epididymal white adipose tissue (WAT) (**Extended Data Figure 1b**). Besides endogenous heme synthesis, cells can also acquire heme from the extracellular environment (i.e., bloodstream). A few transporters have been proposed to cover this function. SLC48A1 appears to have the widest tissue distribution^25–27^, while two other transporters, FLVCR2^28^ and SLC46A1 (a.k.a. HCP1/PCFT)^29^ have restricted expression and likely play important roles in maintaining heme homeostasis in the placenta and the gut, respectively. Intracellular heme increases during adipocyte differentiation^30^, but the relative contribution of endogenous (i.e., synthesis) and exogenous (i.e., import) heme to the total intracellular heme pool in adipocytes has not been determined. To address this question, we evaluated the expression of all the biosynthetic enzymes and putative heme transporters during adipocyte differentiation of primary white, primary brown, and immortalized brown preadipocytes. In addition to detecting robust expression of all heme biosynthetic enzymes in all three cell models, we found that, with the exception of *Cpox*, the expression of all genes increased in a differentiation-dependent manner and that *Alas1* showed the greatest induction (**Extended Data Figure 1c**). Consistent with the erythroid-specific expression, *Alas2* expression was undetectable in adipocytes (**Extended Data Figure 1d**). Of the three putative heme importers, only *Slc48a1* was highly expressed in preadipocytes, with moderately increased levels in fully mature cells (∼2-fold increase) (**Extended Data Figure 1e**). These results suggest that increased endogenous heme biosynthesis, rather than upregulation of heme uptake, is likely responsible for the rise of intracellular heme levels in mature adipocytes.

To assess the dependency of adipocytes on endogenous heme production, we exposed primary brown adipocytes to succinylacetone (SA), a chemical inhibitor of heme biosynthesis, or a media supplemented with heme-depleted serum (HDS). Intracellular heme decreased by a modest ∼15% when cells were grown in heme-depleted media (**Figure 1a**). A much greater reduction (∼60%) was observed when heme biosynthesis was blocked, indicating that the intracellular pool of heme relies primarily on heme synthesis (**Figure 1a**). Consistent with this observation, we found that in response to SA-induced heme deficiency, *Alas1* mRNA and protein were upregulated 4-fold (**Extended Data Figure 2a** and **Figure 1c**). However, since succinylacetone blocks ALAD activity, which is downstream of ALAS1 (**Extended Data Figure 1a**), this does not result in increased heme synthesis. In contrast, depleting exogenous heme in media had minimal to no effects on ALAS1 levels (**Extended Data Figure 2a** and **Figure 1c**). Perhaps surprisingly, inhibition of heme biosynthesis did not impact the ability of cells to develop into mature adipocytes, as the expression of classical markers of differentiated adipocytes (i.e., *Pparγ*, *Fabp4*) and lipid accumulation were comparable across conditions at the end of differentiation (**Extended Data Figure 2a** and **b**). Nevertheless, reduction of intracellular heme levels resulted in a dramatic decrease of *Ucp1* and OXPHOS mRNA and proteins (**Figure 1b** and **c**), indicating that thermogenic potential and mitochondrial function of brown adipocytes are tightly linked to intracellular heme homeostasis.

**Figure 1.**
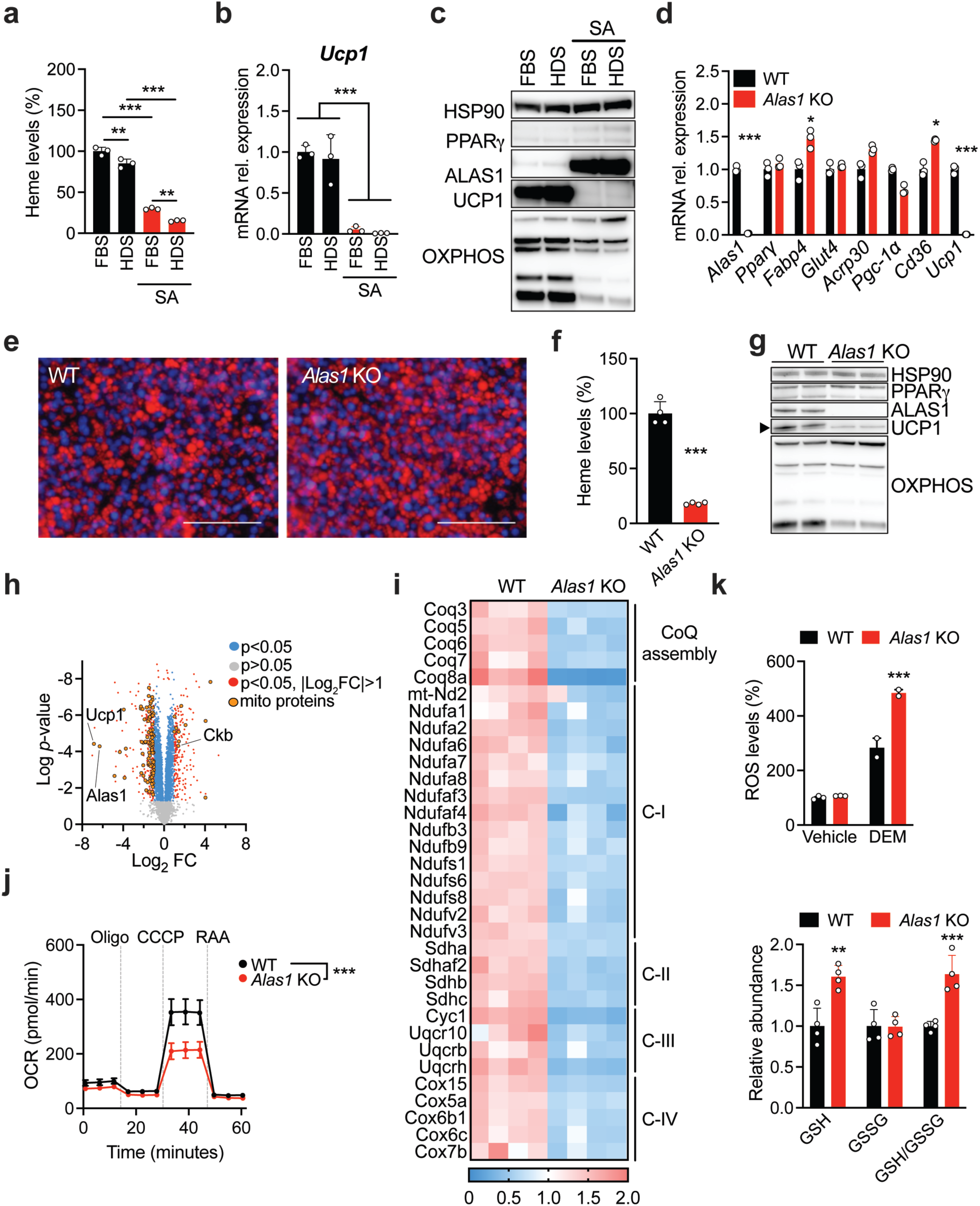
Inhibition of heme biosynthesis induces mitochondrial dysfunction and downregulation of UCP1 in brown adipocytes. **a**) Quantification of intracellular heme levels of primary brown adipocytes differentiated in presence of FBS, heme-depleted serum (HDS), succinylacetone (SA), or HDS and SA. **b**) mRNA relative expression of *Ucp1* in primary brown adipocytes differentiated as described in **a**. **c**) ALAS1, PPARγ, UCP1 and OXPHOS levels in primary brown adipocytes differentiated as described in **a**. **d**) mRNA relative expression of adipogenic and thermogenic genes in WT and *Alas1* KO brown adipocytes. **e**) WT and *Alas1* KO cells show comparable adipogenic differentiation. Lipid accumulation was evaluated on day 8 using the fluorescent dye Nile red (red). Hoechst (blue) was used to stain nuclei. Scale bar = 100 μm. **f**) Quantification of intracellular heme levels in differentiated WT and *Alas1* KO adipocytes. **g**) ALAS1, PPARγ, UCP1 and OXPHOS protein levels in differentiated WT and *Alas1* KO adipocytes. **h**) Volcano plot displaying abundance levels of proteins in *Alas1* KO adipocytes relative to WT. **i**) Relative abundance of components of the electron transport chain complexes detected in **h**. **j**) *Alas1* KO brown adipocytes have impaired mitochondrial respiration. **k**) Hydrogen peroxide levels are significantly increased in *Alas1* KO brown adipocytes upon exposure to diethylmaleate (DEM) for 4 hours. Reduced glutathione (GSH) levels and reduced vs. oxidized GSH (GSH/GSSG) ratio are elevated in *Alas1* KO adipocytes. Data are shown as mean ± SD.* p<0.05, ** p<0.01, ***p<0.001 vs. WT; one-way ANOVA with multiple comparisons and a Tukey’s post-test (a, b, j, k) or two-tailed Student’s *t*-test (d, f).

To confirm the link between heme biosynthesis and the reduction in UCP1, and exclude potential off-target effects of succinylacetone, we generated multiple *Alas1* KO preadipocyte clones using CRISPR (**Extended Data Figure 3a**). Similar to chemical inhibition of heme biosynthesis (**Extended Data Figure 2a**), *Alas1* deletion in immortalized brown preadipocytes did not impact the expression of *Pparγ* and other adipocyte markers (**Figure 1d** and **Extended Data Figure 3b**), resulting in comparable adipocyte differentiation (**Figure 1d** and **e**). However, *Alas1* deletion led to significant reduction of intracellular heme (**Figure 1f**) and, again, downregulation of UCP1 and OXPHOS (**Figure 1d, g** and **Extended Data Figure 3c**). To globally assess the impact of *Alas1* deletion on brown adipocytes, we performed whole-cell proteomics in wildtype and *Alas1* KO cells. We found that ∼30% of significantly reduced proteins in *Alas1* KO cells localize to the mitochondria, including UCP1 and several components of the electron transport chain (**Figure 1h** and **i**). Notably, Creatine Kinase B (CKB), which controls futile creatine cycling in thermogenic fat^31^, was increased in *Alas1* KO cells (**Figure 1h**), suggestive of a compensatory mechanism for the loss of UCP1^31^. A consequence of the extensive reduction of mitochondrial proteins in *Alas1* KO adipocytes was a significant reduction in oxygen consumption rate (OCR) (**Figure 1j**). Besides its critical role in the electron transport chain, heme is also an important cofactor for reactive-oxygen-species-(ROS)-degrading enzymes, such as catalase. Therefore, we measured ROS levels in WT and *Alas1* KO adipocytes. Although we did not detect significant differences between heme synthesis-competent and -deficient cells in basal conditions (**Figure 1k**), we found that glutathione levels, the heme-independent arm of the oxidative stress response, as well as the reduced vs. oxidized GSH/GSSG ratio, were significantly higher in *Alas1* KO cells (**Figure 1k**). However, when exposed to diethylmaleate (DEM), which depletes glutathione in cells, *Alas1* KO adipocytes showed significantly higher levels of ROS compared to WT (**Figure 1k**), indicating reduced, heme-dependent, catalase activity.

To investigate the impact of adipose heme biosynthesis on systemic homeostasis, we generated mice with a specific deletion of *Alas1* in brown adipose tissue (referred to as **BAKO**, **B**AT-specific **A**LAS1 **KO**, *Alas1^fl/fl^*;Ucp1-Cre^32^). To avoid compensation mechanisms, unless noted, all procedures were carried out at thermoneutrality. Unlike whole-body *Alas1* KO mice, which are embryonically lethal^22^, BAKO mice are viable and display normal development under standard housing and feeding conditions (**Extended Data Figure 4a** and **b**). Notably, BAKO BAT showed loss of its distinctive brown color, indicative of reduced heme levels (**Figure 2a** and **b**). Histological analysis also revealed altered morphology, with a striking increase in the size of lipid droplets (**Figure 2c** and **Extended Data Figure 4c**). Despite these differences, energy expenditure was comparable between BAKO and their WT littermates (**Extended Data Figure 4d**), reflecting the marginal role of BAT at thermoneutrality. However, in BAKO mice, activation of adaptive thermogenesis with the β_3_-adrenergic receptor agonist CL316,243 resulted in a significantly blunted response compared to the sustained increase in oxygen consumption and energy expenditure observed in WT mice (**Figure 2d**). Similarly, when exposed to cold (4°C), BAKO mice became rapidly hypothermic, unlike WT mice which were able to maintain their body temperature (**Figure 2e** and **f**). Consistent with the inability of BAKO mice to activate thermogenesis, we observed a marked reduction in UCP1 levels and a dramatic loss of OXPHOS proteins in BAKO BAT (**Figure 2g** and **h**). Transcriptomic analysis of BAT from mice housed at thermoneutrality supported these observations revealing an overall downregulation of genes involved in lipid catabolism, adaptive thermogenesis, and cAMP-mediated signaling (**Figure 2i**, **Extended Data Figure 5a** and **b**), corroborating the whitening of BAKO BAT. On the other hand, to compensate for impaired thermogenesis and lipid catabolism, several biological pathways, including the futile creatine cycle, glutathione-dependent oxidative stress response, and calcium transport, were upregulated in BAKO BAT (**Figure 2i**, **Extended Data Figure 5a** and **b**). These results in BAKO mice align with other models characterized by BAT dysfunction^16,33^ and, although not completely unexpected, reinforce the importance of heme biosynthesis for brown fat function *in vivo*.

**Figure 2.**
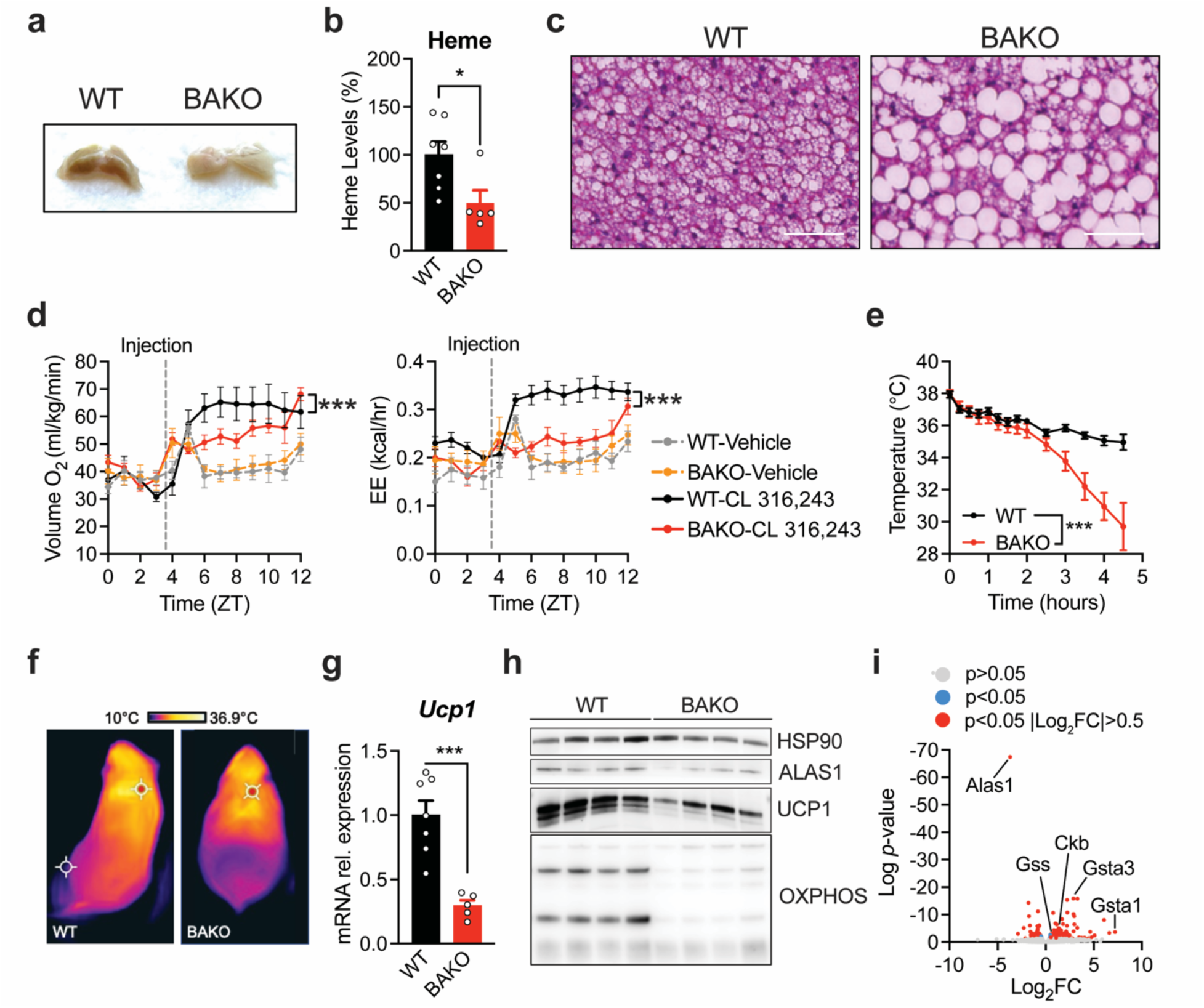
BAT-specific *Alas1* Knockout mice (BAKO) have impaired thermogenesis. **a**) Representative images of BAT appearance in WT and BAKO mice at 30°C after perfusion with PBS-EDTA solution. **b**) BAKO BAT has reduced heme levels (female mice, n = 7 WT, 5 BAKO). **c**) Representative images of hematoxylin and eosin (H&E) staining of BAT depots from WT and BAKO mice maintained at 30°C (female mice, n = 5 per group). Scale bar = 50 μm. **d**) Oxygen consumption and energy expenditure in male WT (n = 6) and BAKO (n = 6) mice housed at 30°C in response to intraperitoneal injection of vehicle (PBS) or the β_3_-adrenoreceptor agonist CL 316,243. **e**) Core body temperature of female WT (n = 9) and BAKO (n = 7) mice under acute cold challenge (4°C). **f**) Forward looking infrared (FLIR) images of WT and BAKO female mice capturing surface body temperature at the 2-hour timepoint of acute cold challenge. **g**) mRNA relative expression of *Ucp1* in BAT of WT and BAKO mice after cold challenge (female mice, n = 7 WT, 5 BAKO). **h**) ALAS1, UCP1 and OXPHOS protein levels in BAT of WT and BAKO mice after cold challenge. **i**) Volcano plot displaying differentially expressed genes (DEGs) in BAT of BAKO mice maintained at 30°C relative to WT BAT. Data are shown as mean ± SEM. * p<0.05, ***p<0.001 vs WT; two-way ANOVA with multiple comparisons and a Tukey’s post-test (e) or two-tailed Student’s *t*-test (b, g).

To understand the extent to which mitochondrial dysfunction and *Ucp1* downregulation in mature brown adipocytes depends on heme depletion, we supplemented both wildtype and *Alas1* KO cells with heme:arginate (HA). Unlike hemin, a form of ferric-chloride-heme adduct that triggers a robust oxidative stress response and is rapidly degraded by cells, HA mimics a slow-release formulation with reduced toxicity, allowing longer exposure to prolonged high levels of treatment^27^. Treatment with HA was sufficient to restore intracellular heme in *Alas1* knockout adipocytes to levels equivalent to those seen in WT cells (**Figure 3a**), indicating that brown adipocytes can indeed uptake exogenous heme. Furthermore, this condition provided an ideal scenario for comparing WT and *Alas1* KO + HA cells, as heme levels are identical, but the heme biosynthesis is either active (in WT) or inactive (in KO). In these conditions, we measured oxygen consumption and found that maximal respiration, induced by CCCP, was completely restored in *Alas1* KO cells supplemented with HA (**Figure 3b** and **c**). However, both basal and uncoupled respiration remained significantly lower in *Alas1* null adipocytes treated with HA compared to WT (**Figure 3b** and **c**). This apparently surprising result can be explained by the effect of HA treatment on *Alas1* KO adipocytes. In fact, HA restored the levels of OXPHOS proteins to a range similar to what is observed in WT adipocytes (**Figure 3d**). However, HA failed to reverse the decrease of UCP1 (**Figure 3d** and **Extended Data Figure 6a**). Consequently, maximal respiration, which relies on the proper function of OXPHOS and is independent of UCP1 (CCCP acts as a chemical uncoupler bypassing UCP1), returned to WT levels. Instead, basal and uncoupled respiration, processes in which UCP1 activity is significantly involved in brown adipocytes, remained significantly lower in *Alas1* KO + HA, as *Ucp1* expression was not restored (**Figure 3d**). Similarly, heme supplementation in primary brown adipocytes treated with SA was unable to revert the SA-induced repression of *Ucp1* (**Figure 3e** and **Extended Data Figure 6b**), despite heme levels being restored (**Extended Data Figure 6c**) as also indicated by return to basal expression of *Alas1* upon HA treatment (**Figure 3e** and **Extended Data Figure 6b**). Thermogenic futile cycles, such as the creatine cycle, also play a role in increasing oxygen consumption in brown adipocytes^3^. Unlike UCP1, these cycles rely on ATP to function. In *Alas1* KO + HA there is an increase in OXPHOS, leading to higher ATP production. This surplus of ATP likely enables futile cycles like the creatine cycle (which is upregulated in *Alas1* KO cells and BAKO mice, **Figure 1h** and **2i**) to function more effectively and may account for the higher uncoupled respiration in *Alas1* KO + HA compared to *Alas1* KO cells (**Figure 3b**), even though the levels of UCP1 remain unchanged between these two conditions (**Figure 3d** and **e**). To comprehensively interrogate the impact of heme biosynthesis inhibition on the transcriptional signature of brown adipocytes, we performed RNA-Seq analysis in primary brown adipocytes treated with SA, HA, or both. Consistent with our previous findings, *Ucp1* was the most downregulated gene in cells exposed to SA, and heme (HA) treatment did not reverse this effect (**Fig. 3f**, left and right panels). Notably, HA alone had minimal influence on transcription, mostly involving genes related to the oxidative stress response (e.g., *Hmox1* upregulation) (**Fig. 3f**, center). Instead, gene ontology enrichment for genes downregulated by SA revealed multiple biological pathways related to energy production, including mitochondrial uncoupling and positive regulation of cold-induced thermogenesis, as well as pathways associated with calcium handling such as host defense and interferon regulation (**Figure 3g**), pathways that have been previously reported to be dysregulated in brown adipocytes lacking UCP1^33^. Based on this evidence, we concluded that brown adipocytes can effectively uptake exogenous heme, and restoration of intracellular heme levels is sufficient to rescue, for the most part, mitochondrial function. However, *Ucp1* downregulation and uncoupled respiration in *Alas1* KO adipocytes are independent of intracellular heme levels.

**Figure 3.**
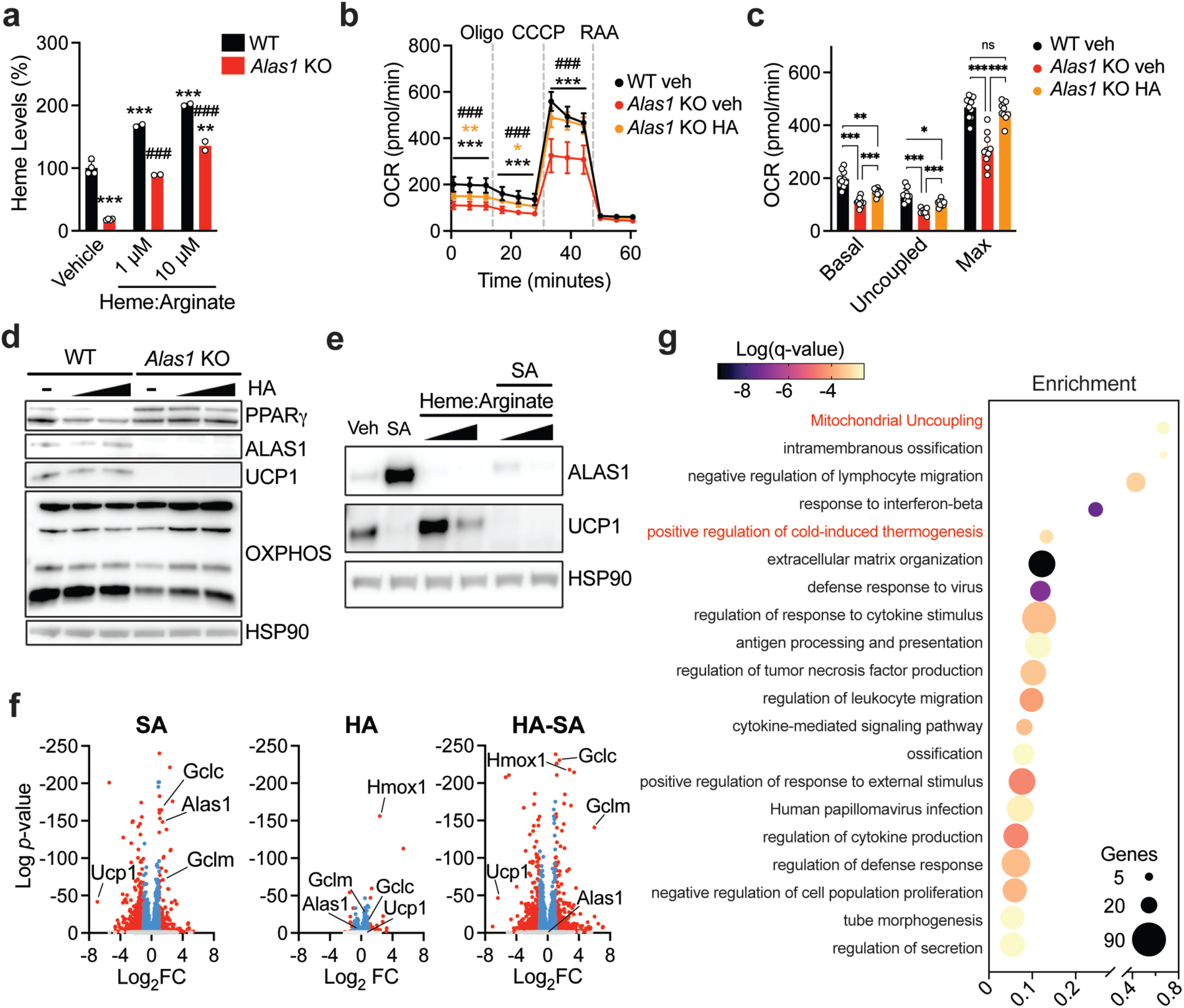
*Ucp1* expression is independent of intracellular heme levels. **a**) Quantification of intracellular heme levels in WT and *Alas1* KO brown adipocytes supplemented with heme-arginate (HA) throughout differentiation. **b**, **c**) Oxygen consumption rates (OCR) of WT and *Alas1* KO brown adipocytes with or without HA. **d**) ALAS1, PPARγ, UCP1, and OXPHOS protein levels in WT and *Alas1* KO brown adipocytes differentiated in the presence of increasing concentration of heme:arginate. **e**) ALAS1 and UCP1 protein levels in primary brown adipocytes treated with heme:arginate, succinylacetone (SA), or both. **f**) Volcano plots of differentially expressed genes (DEGs) in primary brown adipocytes differentiated in the presence of SA, HA or both relative to vehicle-treated cells. Color scale indicates adjusted p-values p>0.05 (grey), adjusted p-values p<0.05 and Log_2_-fold change<|1| (blue), and adjusted p-values p<0.05 and Log_2_-fold change>|1|. **g**) Bubble plot of biological pathways significantly downregulated by succinylacetone. Data are shown as mean ± SD. * p<0.05, ** p<0.01, ***p<0.001 vs WT, ### p<0.001 vs KO-vehicle; one-way ANOVA with multiple comparisons and a Tukey’s post-test (a, b, c).

These unforeseen results led us to hypothesize that the accumulation of ALAS1 substrates, not the depletion of heme, is responsible for the inhibition of *Ucp1* expression. Alas1 produces 5-ALA via condensation of glycine and succinyl-CoA with an 8:8:1 stoichiometric ratio, for 1 molecule of heme requires 8 molecules of glycine and 8 molecules of succinyl-CoA. To gauge how ALAS1 deletion impacts substrate levels, we conducted metabolomic analysis in both WT and *Alas1* KO + HA adipocytes. We found that both glycine and succinyl-CoA, along with their precursors, serine (which generates glycine through serine-glycine one-carbon metabolism) and propionyl-CoA (which is transformed into succinyl-CoA through the reaction catalyzed by the PCCA-MMUT-MCEE complex, **Figure 4e**), were significantly increased in *Alas1* KO adipocytes (**Figure 4a** and **Extended Data Figure 7a**). One of the major sources of propionyl-CoA in cells is the breakdown of the branched-chain amino acids (BCAAs) valine and isoleucine, which also accumulated in *Alas1* KO adipocytes (**Figure 4a** and **Extended Data Figure 7a**). We then asked whether elevated levels of *Alas1* substrates or their precursors could lead to a reduction in *Ucp1* expression in cultured brown adipocytes. To explore this, mature adipocytes were exposed to glycine, serine, valine, isoleucine, propionate, or succinate for 48 hours. Previous research has shown that succinate accumulation does not lead to a decrease in UCP1^34^. In line with these findings, our observations indicated that *Ucp1* expression remained unaffected in brown adipocytes treated with succinate (**Figure 4b** and **c**). However, all the other treatments decreased *Ucp1* (**Figure 4b** and **c**). Notably, only propionate fully phenocopied the effects on *Ucp1* expression seen with the heme synthesis inhibitor succinylacetone (SA) (**Figure 4b** and **c**). This observation led us to inquire about the apparent discrepancy in the impact on *Ucp1* expression when cells are exposed to high levels of valine and isoleucine, whose catabolism generates propionyl-CoA. One explanation is that, under normal conditions, the propionyl-CoA generated from the catabolism of valine and isoleucine is rapidly converted into succinyl-CoA. This metabolite can then be directed into the tricarboxylic acid (TCA) cycle or used for heme biosynthesis. Interestingly, treatment with valine and isoleucine in brown adipocytes significantly elevated intracellular levels of heme (**Figure 4d**), suggesting a potential connection between BCAA homeostasis and heme biosynthesis. To further substantiate the connection between BCAAs and heme, we looked at a comprehensive multi-tissue transcriptomic dataset gathered from a cohort of 500 genetically unique mice (Diversity Outbred, DO500HFD). Strikingly, we discerned a notable correlation between *Alas1* expression and various genes closely linked to BCAA metabolism (**Figure 4f** and **Extended Data Figure 7b**). These genes include Propionyl-CoA carboxylase (*Pcca*), Methylmalonyl-CoA Mutase (*Mmut*), and the recently discovered mitochondrial BCAA transporter *Slc25a44*^17^ (**Extended Data Figure 7b**). Moreover, biological pathway analysis of the top 50 genes whose expression correlates with *Alas1* in adipose tissue revealed an enrichment of genes associated with branched-chain amino acid degradation (**Figure 4f**). Importantly, this correlation was specific to the adipose tissue, as similar analyses conducted in the liver or skeletal muscle did not yield any significant correlations (**Figure 4f** and **Extended Data Figure 7b**).

**Figure 4.**
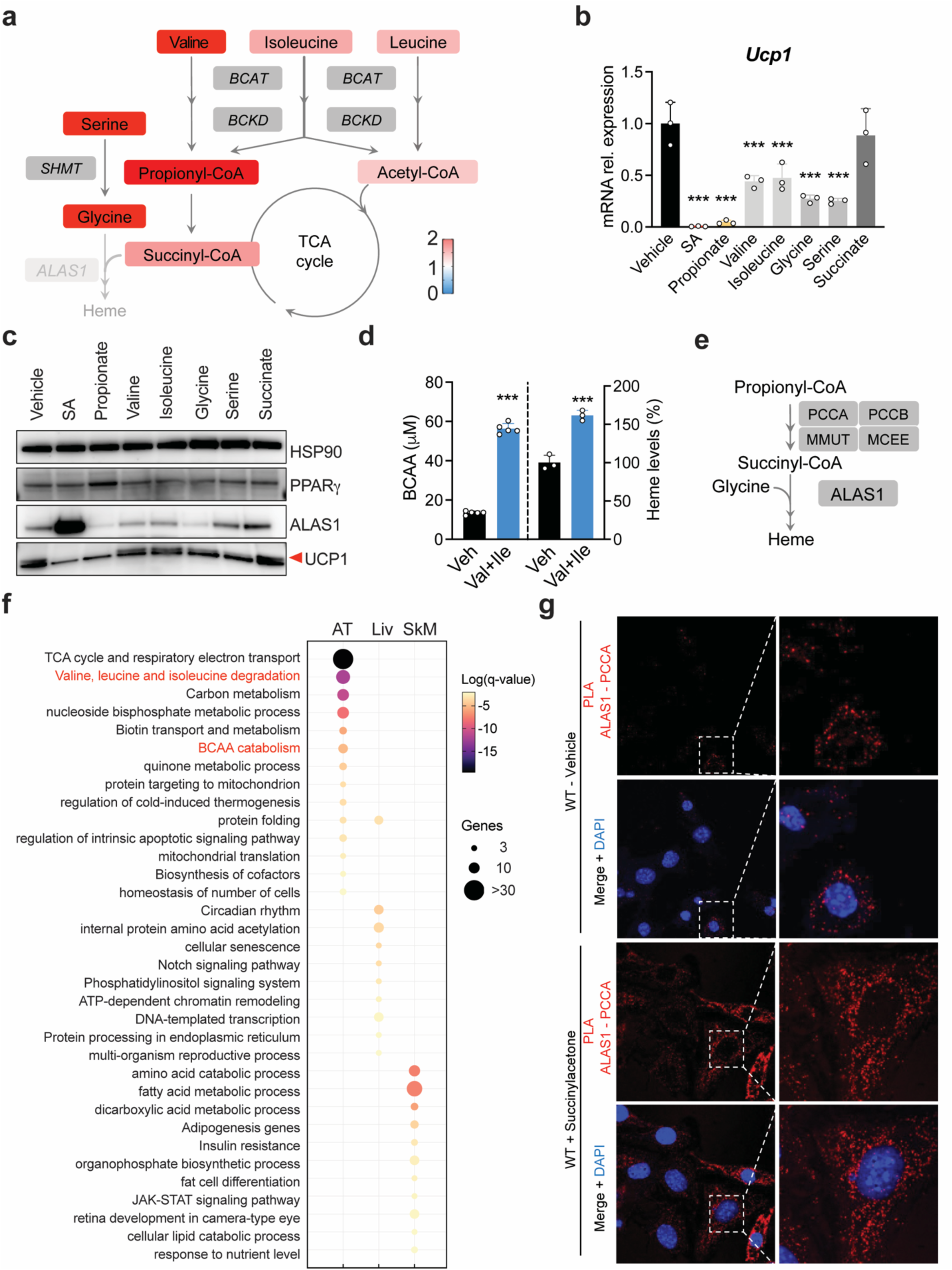
Propionyl-CoA metabolism is linked to heme biosynthesis and *Ucp1* expression in brown adipocytes. **a**) Relative metabolite abundance of glycine- and succinyl-CoA-generating metabolites in *Alas1* KO brown adipocytes relative to WT. **b**) *Ucp1* mRNA levels in primary brown adipocytes treated for 48 hours with glycine- and succinyl-CoA generating metabolites. **c**) ALAS1, PPARγ, and UCP1 protein levels in primary brown adipocytes treated as described in **b**. **d**) Quantification of intracellular BCAA concentrations and heme levels in WT adipocytes differentiated in the presence of 1X or 10X the concentrations of the BCAAs valine and isoleucine. **e**) Schematic of propionyl-CoA conversion to succinyl-CoA. **f**) Bubble plot of biological pathways enriched among the top 50 genes whose expression correlates with *Alas1* in adipose tissue, liver, and skeletal muscle. **g**) Proximity ligation assay of ALAS1 and PCCA in WT brown adipocytes differentiated in the presence or absence of succinylacetone (SA). Punctate foci (red) indicate proximal association of ALAS1 and PCCA. Nuclei counterstain is shown in blue. Data are shown as mean ± SD. ***p<0.001 vs. vehicle; one-way ANOVA with multiple comparisons and a Tukey’s post-test (b) or two-tailed Student’s *t*-test (d).

These observations prompted us to explore the existence of a metabolon complex comprising PCCA and ALAS1, facilitating the conversion of propionyl-CoA into succinyl-CoA which would serve as a readily available substrate for ALAS1, as opposed to using the pool of succinyl-CoA generated by the TCA cycle. Recent research has shown that, in erythrocytes, glutamine is the primary precursor for 5-ALA^35^. Glutamine is converted to α-ketoglutarate and then to succinyl-CoA by α-ketoglutarate dehydrogenase (KDH). Notably, this intermediate does not pass through the TCA cycle because of a direct interaction between the erythroid-specific ALAS2 and KDH^35^. To investigate the hypothesis of an ALAS1-specific metabolon in adipose tissue, we conducted a proximity ligation assay (PLA) in mature brown adipocytes to assess the co-localization of ALAS1 and PCCA within intact cells. Under unstimulated conditions, we observed a discernible yet low-level PLA signal indicative of close proximity between ALAS1 and PCCA (**Figure 4g**). Notably, when ALAS1 levels were induced, as achieved through inhibition of heme biosynthesis by succinylacetone, the signal became significantly greater (**Figure 4g**). Minimal to no PLA signal was detected in *Alas1* KO brown adipocytes, proving the specificity of the PLA signal seen in WT cells (**Extended Data Figure 7c**).

To summarize our findings concisely, we have determined that heme biosynthesis is the primary contributor to intracellular heme levels in brown adipocytes. Inhibition of heme biosynthesis leads to mitochondrial dysfunction and a substantial reduction in *Ucp1* expression. Although supplementing exogenous heme can restore mitochondrial function in heme synthesis-deficient cells, the downregulation of *Ucp1* expression persists even after intracellular heme deficiency is resolved. This persistence does not appear to depend on heme itself but, rather, to the accumulation of the heme precursor propionyl-CoA.

Recently, it was shown that cold exposure promotes uptake of BCAAs in BAT and defects in BCAA catabolism lead to impaired thermogenesis^17^. In a separate study^18^, an extensive analysis of how different tissues use fuels during exposure to cold found that although the contribution of BCAAs to the TCA cycle in BAT and WAT increases in fed mice exposed to cold, it never exceeds 2% of the total TCA cycle metabolism. Our results support a transcriptional and functional link between BCAA catabolism and heme biosynthesis in BAT (**Extended Data Figure 8**). The activation of thermogenesis in BAT is associated with production of mitochondrial ROS, both *in vivo* and *in vitro*^36^. To balance beneficial and detrimental effects on ROS^37^, BAT relies on glutathione and heme-dependent, cold-induced catalases and peroxidases^38^. The assembly of a metabolon including PCCA and ALAS1 suggests that the increased BCAA catabolism in response to cold is channeled into heme production to support mitochondrial function (by providing the cofactor for the electron transport chain) and oxidative stress response. Based on this evidence, the reduced ALAS1 expression in the adipose of obese-diabetic subjects^24^, and the concurrent elevation in BCAAs and propionate levels^39–46^, we posit that heme biosynthesis in adipocytes may serve as an important metabolic sink for BCAA clearance. Further studies are deemed to quantitively assess the contribution of BCAAs to heme biosynthesis and understand the molecular mechanisms underlying propionyl-CoA-dependent repression of *Ucp1* expression. However, our work integrates current literature by providing a model for how brown fat utilizes BCAAs during stimulated conditions.

## ACKNOWLEDGEMENTS

We thank Dr. Dudley Lamming, Dr. Caroline Alexander, Dr. Judith Simcox, and Helaina Von Bank for critical input and reagents; the University of Wisconsin-Madison Genome Editing and Animal Model core for assistance with the generation of the *Alas1* floxed mouse model; Dr. Nadia Rosenthal at The Jackson Laboratory for sharing the RNA-Seq data of the Diversity Outbred mice. This work was supported by the National Institute of Health grants 1R35GM150899 (AG), P41GM108538 (JJC), R35GM147014 (JF), the Wisconsin Partnership Program at the University of Wisconsin School of Medicine and Public Health grant WPP5451 (AG), the DRC at Washington University P30 DK020579 (AG). DJD is supported by the National Institute On Aging training grant T32AG000213, HB is supported by NIH NCATS awards to UW-ICTR TL1TR002375 and UL1TR002373.

## AUTHOR CONTRIBUTIONS

DJD and AG conceived the project, designed research, and analyzed data. DJD, JKH, and MF performed in vivo experiments. DJD, JKH, NC, and YB carried out cell-based assays. DJD and HB performed RNA-Seq analyses. SJ performed metabolomic analyses. AJ, KO, and ES prepared samples and carried out proteomics studies. MK and AG performed correlation expression analysis. RAA, VLC, ADA, JJC, and JF provided advice and reagents. DJD and AG wrote the manuscript and integrated comments from the other authors.

## DATA AVAILABILITY

Source data table for Figures 1-4, Extended Data Figures 1-7, full scans of all western blots, proteomics and transcriptomics data, and supplementary tables will be available at the time of publication.

## METHODS

### Reagents

Oligomycin A (#75351), Carbonyl cyanide 3-chlorophenylhydrazone (CCCP; #C2920), rotenone (#R8875), antimycin A (#A8674), 3-isobutyl-1-methylxanthine (IBMX; #I5879), triiodothyronine (#T5516), insulin (#I6634), dexamethasone (#D1159), BSA (#A7030), L-arginine (#A8094), L-valine (#V0500), L-isoleucine (#I2752), L-glycine (#G7126), L-serine (#S4311), succinate (#S9637), propionate (#P5436), succinylacetone (#D1415), puromycin (#P8833), CL 316,243 (#C5976), Tween-20 (#P1379) and sodium deoxycholate (#D6750) were purchased from Sigma-Aldrich. Rosiglitazone (#71740) and hemin chloride (#16487) were purchased from Cayman Chemical. Complete EDTA-free protease inhibitor cocktail (#11836170001) was obtained from Roche. DMEM and other Gibco-branded cell culture products were purchased from Thermo Fisher. Heme-depleted FBS was prepared by treating FBS with 20 mM ascorbic acid (Sigma Aldrich #A2218) for 16 hr, followed by 48 hr dialysis against PBS. Heme depletion was verified by measuring optical absorbance at 405 nm. gRNAs, primers and probes were purchased from IDT. Collagenase (#LS004196) was purchased from Worthington Biochemical. TMB (#TMBW010001) was purchased from Surmodics. West Pico ECL (#34577) was purchased from Pierce. DC assay (#5000116) was purchased from BioRad. Cells were routinely tested for mycoplasma and were never positive. Primary antibodies used in this work are: HSP90 (GeneTex #101423), ALAS1 and PPARγ (Santa Cruz #sc137093 and #sc7273), UCP1 (Abcam #10157682, Thermo Fisher #PA124894), OXPHOS (Invitrogen #458099) and PCCA (Proteintech #219881AP). Secondary antibodies used are goat anti-mouse and mouse anti-rabbit (Jackson Immunoresearch #115035146 and #211032171). Antibody dilutions for western blot and PLA will be available at the time of publication.

### Brown adipocyte culture

Primary brown adipocytes were isolated and cultured as described previously^47^. Immortalized brown preadipocytes were kindly provided by Dr. Judith Simcox. Cells were maintained in DMEM with 25mM glucose (Thermo Fisher #11965118), 10% FBS (Gemini #100106), 10mM HEPES (Thermo Fisher #15630080), 1mM sodium pyruvate (Thermo Fisher #11360070), GlutaMax (Thermo Fisher #35050061) and Pen/Strep (Thermo Fisher #15240062). Upon reaching confluency, cells were treated with a differentiation cocktail for 2 days, composed of insulin (5 μg/mL), triiodothyronine (T3; 1 nM), dexamethasone (500 nM), 3-isobutyl-1-methylxanthine (IBMX; 500 μM) and rosiglitazone (1 μM). After 2 days, media was switched to differentiation maintenance media composed of insulin (5 μg/mL) and T3 (1 nM) and replaced every 2 days until cells became mature adipocytes. Immortalized brown adipocytes were harvested at day 8 for all experiments, and primary brown adipocytes were harvested at day 6 unless noted otherwise. For relevant experiments, heme-arginate (HA) solution was prepared as a 1:10 molar ratio, with 10 mM hemin chloride + 100 mM L-arginine in a 50/50 v/v solution of 0.2 M KOH and 100% ethanol. HA supplementation was performed throughout differentiation with media changes at a final concentration of 10 μM. Fluorescence microscopy to visualize lipid droplets was performed on mature adipocytes using AdipoRed (1:250 dilution, Lonza #PT-7009) and Hoechst 33342 (1 μg/mL, Thermo Fisher #62249) in DMEM for 15 minutes at 37°C. Cells were washed twice with PBS prior to imaging.

### Generation of Alas1^-/-^ preadipocytes

Alas1 gRNAs were cloned into the backbone of PX459 plasmid backbone as described previously^48^. pSpCas9(BB)-2A-Puro (PX459) V2.0 was a gift from Feng Zhang (Addgene plasmid # 62988; http://n2t.net/addgene:62988; RRID:Addgene_62988). Immortalized brown preadipocytes were transfected with PX459 harboring Alas1 gRNAs using PEI at a 3:1 PEI:DNA mass ratio. gRNA sequences will be available at the time of publication. Two days after transfection, cells were selected with puromycin (2 μg/mL) for six days. Puromycin-selected cells were plated at low density to generate monoclonal colonies, which were then screened by PCR genotyping, western blotting, and qPCR to confirm genetic knockout of *Alas1*. Primer sequences will be available at the time of publication.

### Mitochondrial bioenergetics measurements

Oxygen consumption rate (OCR) was quantified using an Agilent Seahorse XF^e^96 instrument as previously described^16^. Briefly, WT and Alas1 KO brown adipocytes were re-plated onto gelatin-coated XF^e^96 plates at day 8 of differentiation at a density of 30,000 cells/well. Two days after re-plating, cells were equilibrated in serum-free DMEM (Sigma Aldrich #D5030) supplemented with 25 mM glucose (Sigma Aldrich #G8769), 10 mM sodium pyruvate, 2 mM L-glutamine (Thermo Fisher #25030081) and 5 mM HEPES (final pH = 7.40) for 1 hour prior to assay initiation. The mitochondrial stress test protocol followed 3-minute mixing cycles followed by 2-minute measurement cycles for 3 measurements. Basal respiration was measured first, followed by injections to achieve final concentrations of 1 μM oligomycin, 2 μM CCCP, and 2 μM rotenone + Antimycin A, respectively.

### Quantitative PCR and RNAseq

Total RNA was extracted from cells or tissues in TRIzol (Thermo Fisher # 15596018), purified using the Direct-Zol RNA MiniPrep plus kit (Zymo Research #R2052) and quantified using a Take3 plate with a Synergy H1 plate reader (BioTek). Taqman-based quantitative real time PCR was performed using the SuperScript III Platinum One-Step qRT-PCR reagent (Thermo Fisher #11732088). 20 ng RNA was loaded per reaction in a 10 μL volume, with samples run in triplicate and multiplexed to normalize to the internal housekeeping control gene 36B4. qPCR protocol and subsequent analysis was executed using a QuantStudio 5 thermal cycler (Applied Biosystems). Primer and probe sequences will be available at the time of publication. For RNAseq, total RNA was extracted as described above. 500 ng RNA was used for library preparation. mRNA was purified using poly-T oligo-attached magnetic beads, then utilized for cDNA synthesis library prep followed by end repair, A-tailing, adapter ligation, size selection, amplification, and purification. Libraries were sequenced using an Illumina X plus platform using 150 base pair reads at a depth of ∼20 million reads per library. Reads were mapped to the mouse reference genome (GRCm39) using Kallisto. Reads aligning uniquely to a single gene were used in downstream analysis. Gene expression values were calculated using DESeq2. Differentially expressed genes (DEGs) were calculated with 4 replicates per condition at an adjusted p-value of p<0.05 and were subsequently used for pathway analysis using Metascape^49^. Heatmaps were generated using RStudio (package ‘pheatmap’).

### BCAA quantification

Quantification of intracellular BCAAs was performed using the BCAA-Glo^TM^ Assay Kit (Promega #CS306201) as per manufacturer’s instructions. Briefly, WT and Alas1 KO brown adipocytes were differentiated in media supplemented with vehicle or L-valine and L-isoleucine to achieve a concentration 10 times higher than the concentrations found in DMEM. On day 8 of differentiation, cells were lysed and 50 μL lysate was transferred to a new 96-well plate followed by the addition of detection reagent and incubation at room temperature for 90 minutes. Luminescence was measured with a Syngery H1 plate reader (BioTek) and BCAA levels were interpolated based on a L-leucine standard curve.

### Reactive oxygen species (ROS) quantification

Production of hydrogen peroxide (H_2_O_2_) in WT and Alas1 KO brown adipocytes was measured using the ROS-Glo^TM^ H_2_O_2_ assay kit (Promega #G8820) as per manufacturer’s instructions. The assay was performed on day 9 of differentiation in 96-well format. 80 μL of fresh media were added to each well. Cells were then incubated for 2 hours at 37°C. After incubation, 20 μL of H_2_O_2_ substrate solution with or without 5 mM diethylmaleate (DEM) was added to each well, and cells were incubated again for 4 hours at 37°C. Then, 100 μL of ROS-Glo^TM^ detection solution was added to each well and incubated for 20 minutes at room temperature before measuring luminescence signal with a Synergy H1 plate reader (BioTek).

### Protein quantification and Western blotting

Cell and tissue protein lysates were prepared in 0.5% sodium deoxycholate solution and mechanically disrupted via sonication (5 pulses, 30% amplitude, 3s/2s on/off) on ice. Protein quantification was performed using DC assay via interpolation with a BSA standard curve. For western blot, samples were loaded onto 4-12% bis tris gels (Thermo Fisher #NW04120) and separated by electrophoresis at 165 volts for 60-90 minutes. Proteins were transferred to nitrocellulose membranes using the iBlot2 system (Invitrogen #IB21001), then treated with blocking buffer (5% BSA in TBS-Tween 0.1%) for 1 hour at room temperature. Membranes were incubated overnight at 4°C in primary antibody diluted in blocking buffer, washed three times for 10 minutes with TBS-Tween 0.1%, then incubated in HRP-conjugated secondary antibody diluted in blocking buffer for 1 hour at room temperature. Membranes were washed again in TBS-Tween 0.1% as described above, and signal was revealed with Pico ECL reagent imaged using a ImageQuant 800 (Amersham).

### Quantification of heme

Enzymatic detection of heme was performed as previously reported^16^. Briefly, apoperoxidase (ApoHRP; Calzyme Laboratories #239A0000) was extracted with acidic acetone for 15 minutes at room temperature to remove residual heme, then centrifuged at 2,000 x g for 5 minutes at 4°C to pellet. Overlying acidic acetone was removed, the pellet was air dryed for 5 minutes and reconstituted in water. 10 μg of protein lysates in 90 μL PBS were added to 10 μL of 50 μM ApoHRP and incubated for 5 minutes at RT. 10 μL of ApoHRP-protein solution were transferred in duplicate to clear bottom 96-well plates, and 200 μL TMB substrate were added to each sample. Heme-reconstituted HRP activity was measured at 652 nm using a Synergy H1 plate reader (BioTek) and heme content was interpolated using a hemin-ApoHRP standard curve.

### Metabolomics

Mature adipocytes were washed twice with ice cold PBS and lysed via addition of LC/MS-grade 80:20 methanol:H_2_O (Fisher #A4561; Sigma Aldrich #900682) at -80°C for 15 minutes. Metabolite extracts were transferred to 1.5 mL tubes, vortexed, then centrifuged at 21,300 x g at 4°C for 15 minutes. Supernatant was transferred to a new tube and held on ice, while the protein pellet was extracted again via resuspension in LC/MS-grade 80:20 methanol:H_2_O and spun again. Both extracts were pooled, dried down under nitrogen stream, and reconstituted in LC/MS-grade H_2_O. Samples were analyzed using a Thermo Q-Exactive mass spectrometer coupled to a Vanquish Horizon UHPLC. Samples were separated on a 100 x 2.1 mm, 1.7 μM Acquity UPLC BEH C18 Column with a gradient of solvent A (97:3 H_2_O:methanol, 10 mM TBA (Sigma Aldrich #90781), 9 mM acetate (Fisher #11350), pH = 8.2) and solvent B (100% Methanol). The gradient utilized is: 0 min, 5% B; 2.5 min, 5% B; 17 min, 95% B; 21 min, 95% B; 21.5 min, 5% B. The flow rate utilized is 0.2 mL/min with a 30°C column temperature. Data were collected on a full scan negative mode at a resolution of 70K. Settings for the ion source were: 10 aux gas flow rate, 35 sheath gas flow rate, 2 sweep gas flow rate, 3.2 kV spray voltage, 320°C capillary temperature and 300°C heater temperature. Metabolites were identified based on exact m/z ratios and retention times determined with standards. Metabolite levels were normalized to total protein content, and data was analyzed using MAVEN software^50^.

### Proteomics

Cell pellets were resuspended in denaturing buffer composed of 5.4 M guanidine hydrochloride (Sigma Aldrich #G4505) in 100 mM Tris (Life Technologies #15568025), pH = 8.0 and incubated at 100°C for 10 minutes. Samples were diluted with LCMS grade pure methanol to bring samples to 90% methanol v/v and centrifuged at 9,000 x g for 5 minutes to pellet proteins. Overlying supernatant was removed and samples dried for 5 minutes at room temperature, resuspended in lysis buffer composed of 8 M urea (Fisher #U15500), 100 mM Tris, 10 mM TCEP (Sigma Aldrich #75259), 40 mM chloroacetamide (Sigma Aldrich #22790), and then vortexed. Proteins were then digested overnight with 2 μg LysC (Wako #12902541) at room temperature on a shaker. The following morning, LysC digestion was quenched by addition of 100 mM Tris to bring urea concentration to 2 M, and proteins were further digested by the addition of 2 μg trypsin (Promega #V5113) for 4 hours at room temperature on a shaker. Trypsin digestion was quenched by the addition of ∼20 μL LCMS grade 10% TFA (Thermo Fisher #85183) to acidify the sample to a pH between 1-2. Samples were further centrifuged at 9,000 x g for 5 minutes. Desalting was performed using Strata X columns, which were equilibrated with LCMS-grade 100% acetonitrile (Fisher #A9551) and LCMS-grade 0.2% formic acid (Thermo Fisher #85178) over two rounds. Peptides were loaded onto the column, washed with 0.2% formic acid, and eluted in elution buffer composed of 80% acetonitrile and 0.2% TFA, then dried down in a vacuum centrifuge. On day of injection, samples are reconstituted in 0.2% formic acid to a final concentration of 1-2 μg/μL and injected onto the LC-MS.

### Proximity ligation assay (PLA)

PLA was utilized to assess in situ protein-protein interactions as previously described^51^. Briefly, adipocytes were differentiated for 8 days and re-plated onto glass coverslips coated with 0.1% gelatin solution for two additional days. Adipocytes were then fixed with 4% paraformaldehyde (Santa Cruz #sc-281692) for 20 minutes at room temperature, washed three times with PBS, permeabilized with 0.3% Triton-X-100 (Fisher #BP151) for 15 minutes, and washed three times with PBS. Samples were blocked in 1% BSA (in PBS) for 1 hour at room temperature and processed for PLA using the DuoLink in situ Red Starter Kit (Millipore Sigma #DUO92101) per manufacturer’s instructions. After washing three times with PBST, slides were mounted with Prolong^TM^ Glass Antifade Mountant with NucBlue^TM^ Stain (Thermo Fisher #P36985). PLA signals were detected as discrete punctate foci using a Leica SP8 confocal microscope to provide intracellular localization analysis of protein interactions. PLA foci were quantified using ImageJ.

### Mouse studies

All animal work procedures were approved by the institutional animal care and use committee of the University of Wisconsin-Madison School of Medicine and Public Health and conducted in accordance with ethical regulations and policies. *Alas1*-floxed mice within the C57BL6/J background were generated by the Genome Editing and Animal Model (GEAM) core at UW-Madison, harboring LoxP sites flanking exon 4 and 5 of the *Alas1* gene. To generate brown adipose specific *Alas1* knockout mice (BAKO), *Alas1*^flox/flox^ mice were crossed with the Ucp1-Cre strain^32^ (Jax, strain# 024670). Floxed littermates negative for the Cre transgene were used as wild-type controls for all studies. No morphological, behavioral or locomotor abnormalities were observed in BAKO mice. Animals were born and housed at room temperature with a 12-hour light/dark cycle, then transferred to thermoneutrality (30°C) for at least two weeks prior to experiments. Animals were fed standard chow ad libitum (Teklad, diet #2019). For molecular analysis of tissues, mice were euthanized via CO_2_ inhalation at around ZT5 and perfused with ice-cold PBS supplemented with 10 mM EDTA.

### Response to acute cold exposure

To measure core body temperature, temperature transponders (BMDS #IPTT-300) were implanted via intraperitoneal delivery under isoflurane anesthesia and read using a RFID reader (BMDS #DAS-8027-DLU) at least one week prior to cold challenge. Mice were individually caged with free access to water and minimal bedding following transfer to 4°C at around ZT3. Core body temperature measurements were recorded every 15 or 30 minutes for up to 5 hours. Mice whose body temperature dropped below 31°C were considered hypothermic and euthanized.

### Energy balance studies

Measurements of energy balance (VO_2_, VCO_2_, RER, heat production, activity) were determined in a controlled environment using a computer-controlled open circuit system (Oxymax) integrated into a Comprehensive Laboratory Animal Monitoring System (Columbus Instruments) set to a 12-hour light/dark cycle at thermoneutrality (30°C) as previously described^16^. To test the response of animals to β_3_-adrenergic stimulation, animals were acclimated to metabolic cages for one week, then received intraperitoneal injections of PBS or CL 316,243 (1 mg/kg) at ZT4.

### Histology

Following euthanasia, brown adipose tissue was collected from animals and fixed in 10% neutral buffered formalin for 24 hours, extensively washed with PBS (pH = 7.4) and stored in 70% ethanol. Tissues were dehydrated, embedded in paraffin, and 3 mm thick sections stained with hematoxylin and eosin by the UW Experimental Animal Pathology Lab. Images were acquired using an EVOS M5000 Imaging System (Thermo Fisher). Lipid droplet size was quantified using ImageJ software.

### Statistics

Results from *in vitro* assays and cell culture data are presented as mean ± s.d. Data generated in mouse studies are presented as mean ± s.e.m. The number of mice used in each experiment is indicated in the figure legends. Statistical analysis was performed on Prism software (GraphPad) using Student’s t-test for comparisons between two groups, one-way ANOVA with multiple comparisons for assessment of more than two groups, and two-way ANOVA with multiple comparisons for repeated measurements. Comparisons among specific groups were done using post-tests as indicated in the figure legends.

---

**Extended Data Figure 1:**
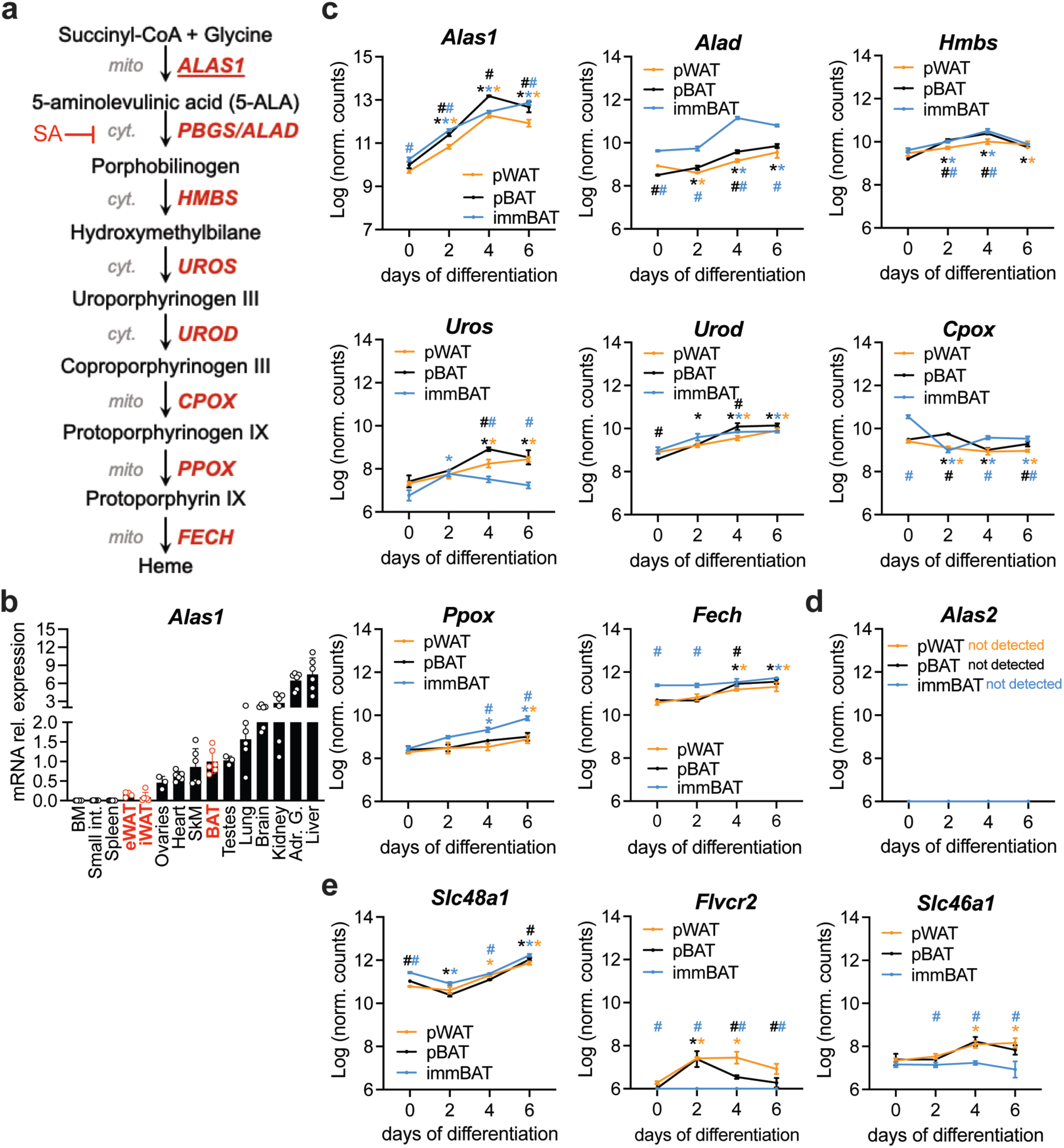
Expression analysis of the heme biosynthesis and uptake pathways in differentiating adipocytes and mouse tissues. **a**) Schematic of heme biosynthesis, an eight-step pathway distributed between the mitochondria and cytosol. 8 molecules of succinyl-CoA and glycine are required to produce 1 molecule of heme. Substrates (black), enzymes (red) and subcellular compartment (gray). Succinylacetone (SA) chemically inhibits ALAD/PBGS enzymatic function. **b**) mRNA relative expression of Alas1 in various mouse tissues collected from wild type 18-week-old male (n = 3) and female (n = 3) mice. **c**) mRNA counts of genes within the heme biosynthesis pathway in primary white (yellow), brown (black) or immortalized brown (blue) adipocytes throughout differentiation. **d**) mRNA counts of erythrocyte-specific ALA synthase (Alas2) was not detected in any adipocyte lines. **e**) mRNA counts of putative heme transporters in primary white (yellow), brown (black) or immortalized brown (blue) adipocytes throughout differentiation. Data shown as mean ± SD. *, *, * p<0.05 vs day 0 of differentiation of pBAT, pWAT, or immBAT. #, # p<0.05 vs same time point of pWAT; one-way ANOVA with multiple comparisons and a Tukey’s post-test.

**Extended Data Figure 2:**
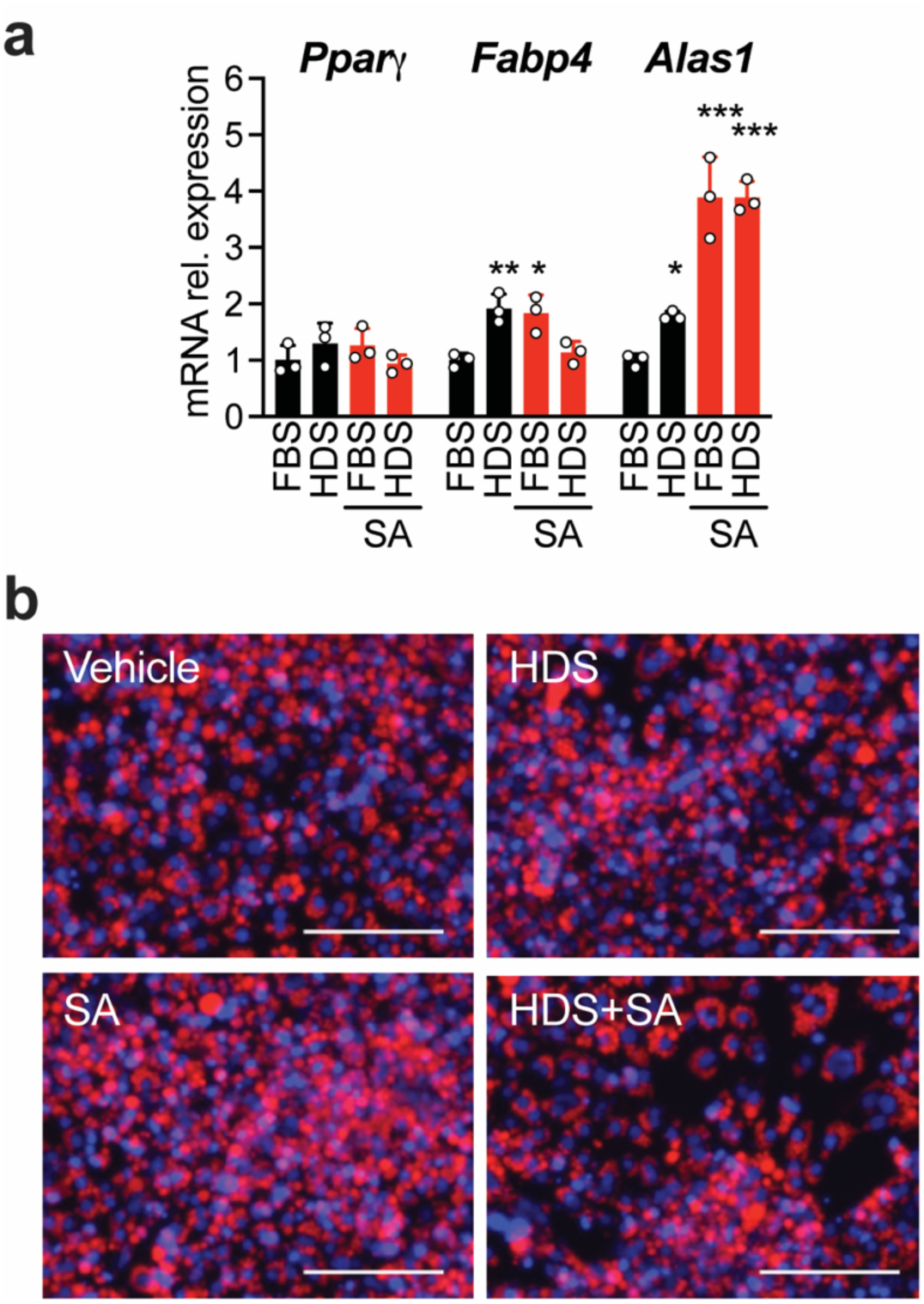
Alteration of heme homeostasis does not impact adipocyte differentiation. **a**) mRNA levels of adipogenic markers (*Pparγ* and *Fabp4*) and *Alas1* in primary brown adipocyte differentiated in the presence of FBS or HDS with or without succinylacetone. **b**) Fluorescence microscopy images of primary brown adipocytes differentiated as in **a**. Lipid droplets were stained using the fluorescent dye Nile red (red), while Hoechst was used for nuclei counterstain (blue). Scale bar = 150 μm. Data shown as mean ± SD. * p<0.05, ** p<0.01, ***p<0.001 vs. FBS; one-way ANOVA with multiple comparisons and a Tukey’s post-test.

**Extended Data Figure 3.**
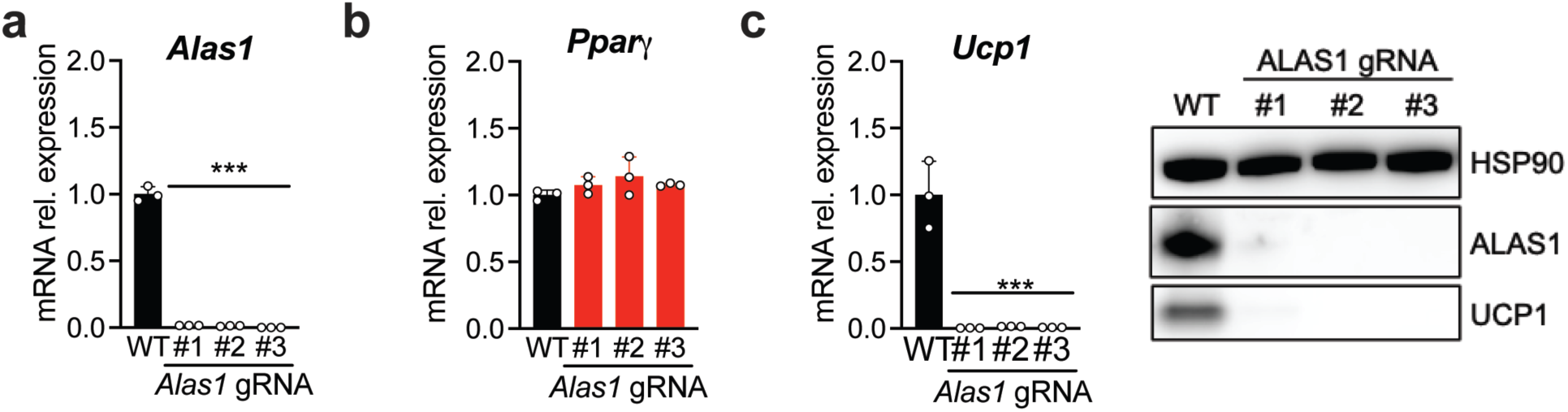
Effect of *Alas1* deletion in brown preadipocytes. **a**) mRNA levels of *Alas1* in three representative *Alas1* KO clones generated using three distinct gRNAs. **b**) *Pparγ* expression is not impacted by ALAS1 deletion. **c**) UCP1 levels are significantly reduced in all *Alas1* KO clones. Data are shown as mean *** p<0.001 vs. WT. Data shown as mean ± SD. * p<0.05, ** p<0.01, ***p<0.001 vs. WT; one-way ANOVA with multiple comparisons and a Tukey’s post-test.

**Extended Data Figure 4.**
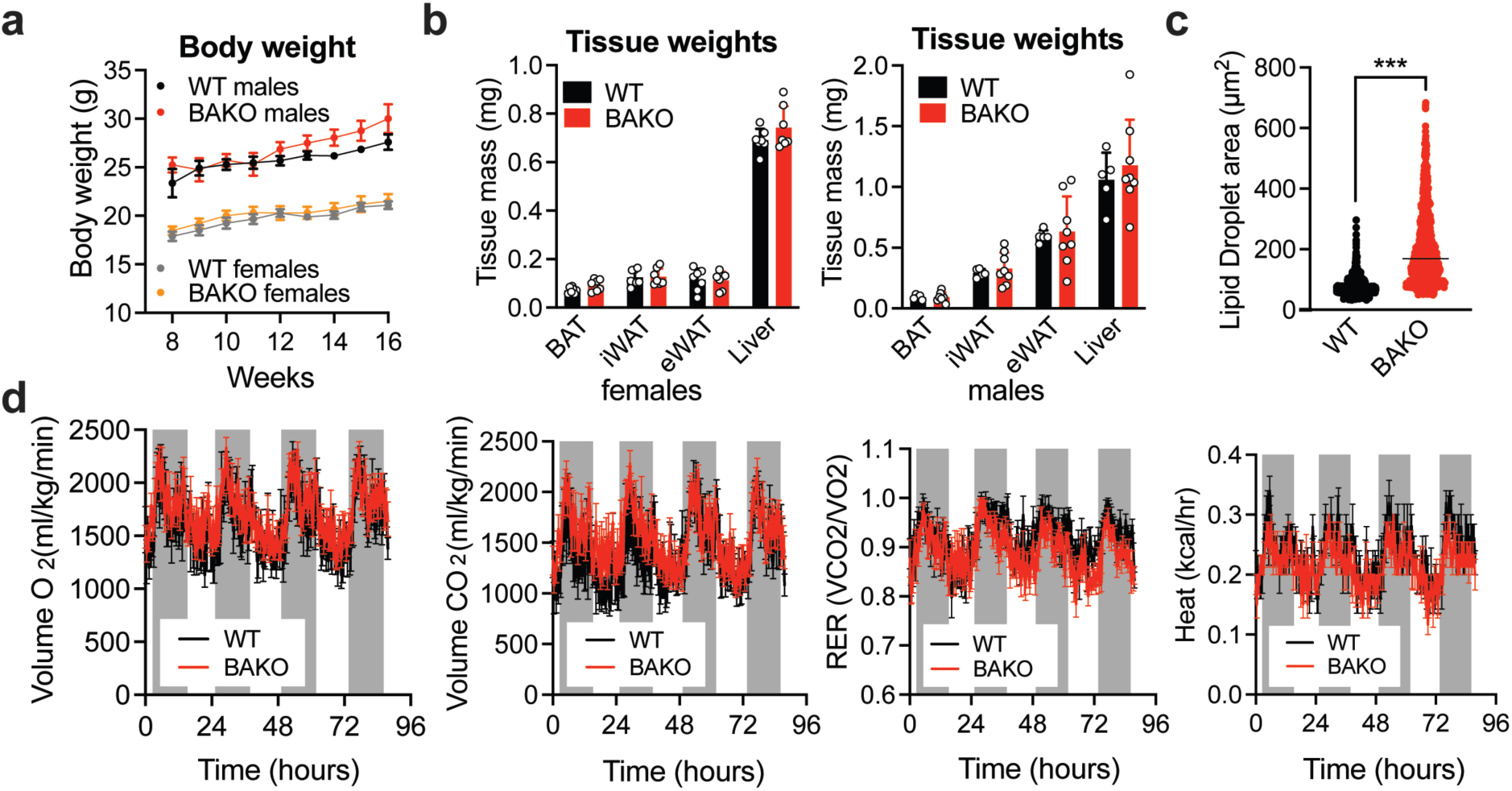
BAKO mice display normal development and energy expenditure at thermoneutrality. **a**) Body weights of WT (male n = 5; female n = 9) and BAKO (male n = 8; female n = 8) mice from 8 to 16 weeks of age. **b**) Tissue weights of BAT, inguinal WAT (iWAT), epididymal WAT (eWAT) and liver from WT (male n = 5; female n = 9) and BAKO (male n = 8; female n = 8). **c**) Measurements of lipid droplet size in BAT collected from female WT and BAKO BAT housed at thermoneutrality. **d**) VO_2_, VCO_2_, respiratory exchange ratio (RER) and heat production of 12-week-old WT (n = 5) and BAKO (n = 7) male mice at thermoneutrality. Data are show as mean ± SEM. *** p<0.001 vs. WT; two-tailed Student’s *t*-test.

**Extended Data Figure 5.**
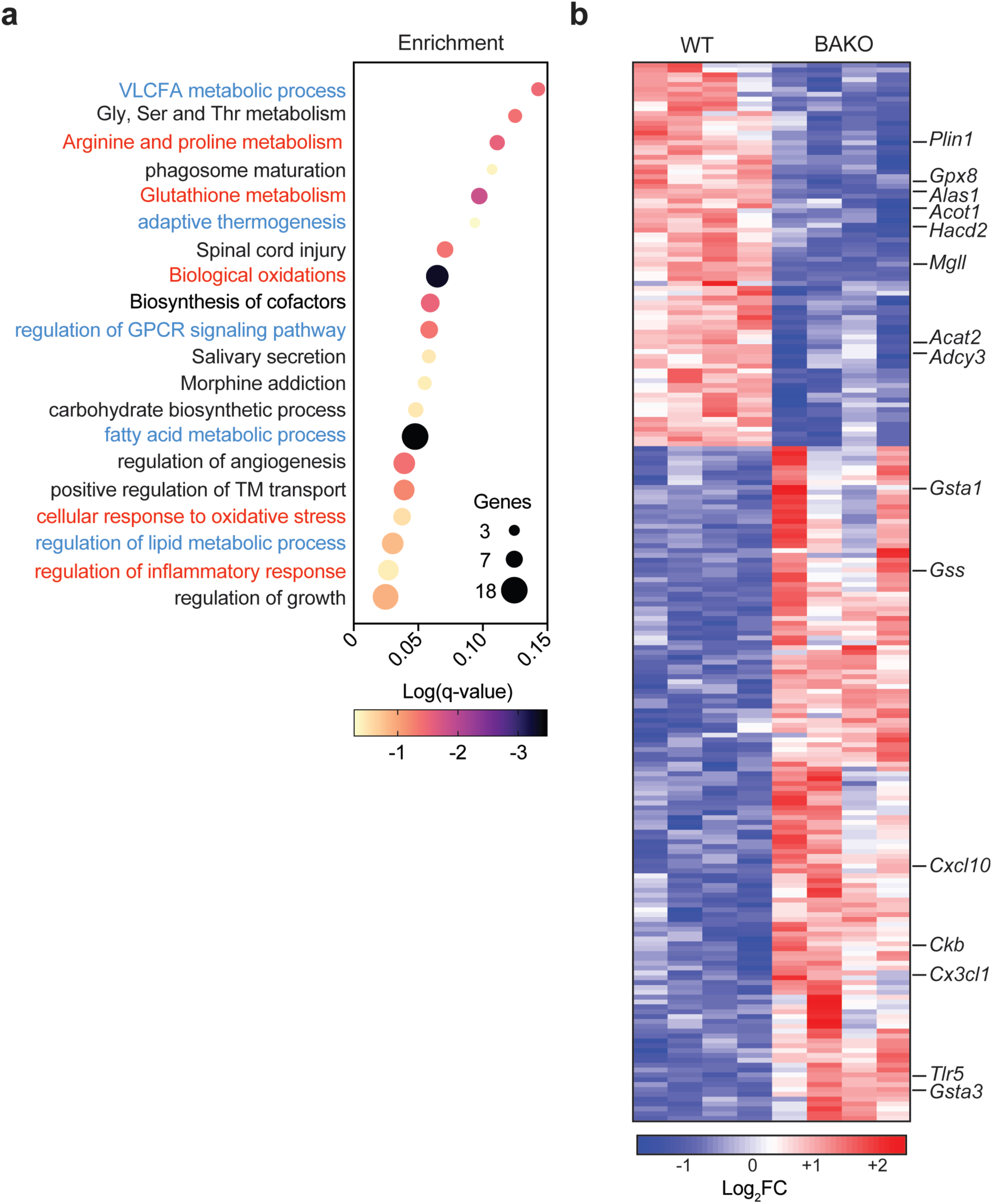
Impact of *Alas1* deletion on the transcriptional signature of BAT. **a**) Biological pathways (BP) differentially enriched between WT and BAKO BAT. BP relative to lipid metabolism, GPCR signaling, thermogenesis, and calcium homeostasis, are reported in blue (downregulated) or red (upregulated). **b**) Heatmap of DEGs in WT and BAKO BAT, with highlighted genes involved in adaptive thermogenesis, oxidative stress response, fatty acid, amino acid, TCA, and glutathione metabolic processes.

**Extended Data Figure 6.**
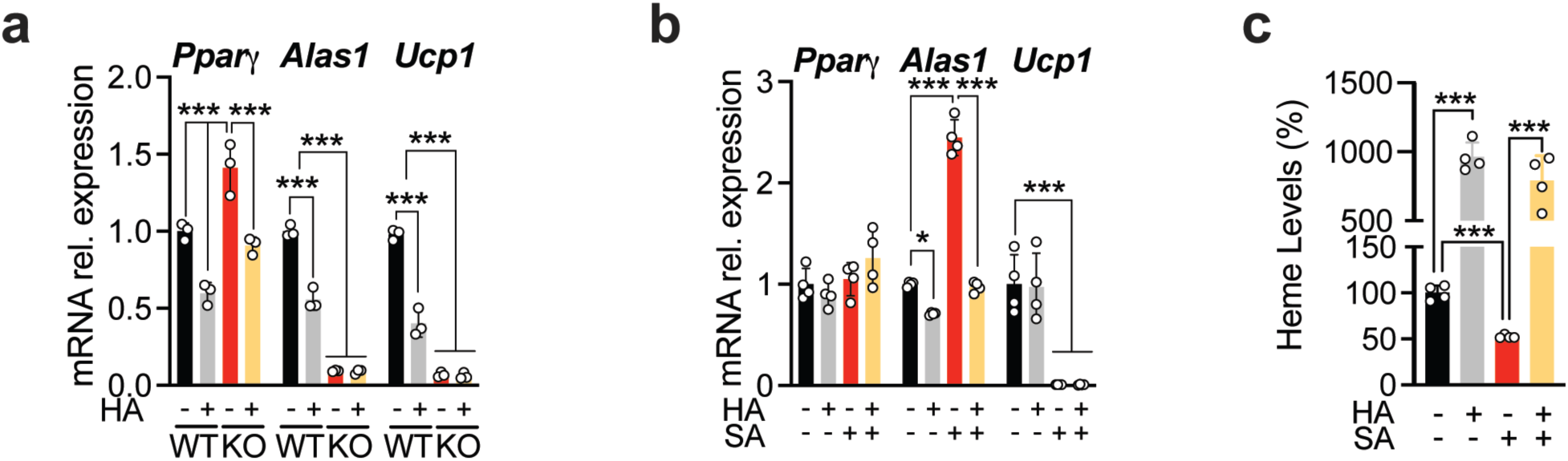
Heme supplementation does not rescue *Ucp1* expression in heme synthesis-deficient brown adipocytes. **a**) *Pparγ*, *Alas1*, and *Ucp1* mRNA levels of WT and *Alas1* KO adipocytes treated with HA throughout differentiation. **b**) *Pparγ*, *Alas1*, and *Ucp1* mRNA levels of primary brown adipocytes treated with SA, HA, or both throughout differentiation. **c**) Heme levels in primary brown adipocytes treated with SA, HA, or both. Data are show as mean ± SD. * p<0.05, *** p<0.001; one-way ANOVA with multiple comparisons and a Tukey’s post-test.

**Extended Data Figure 7.**
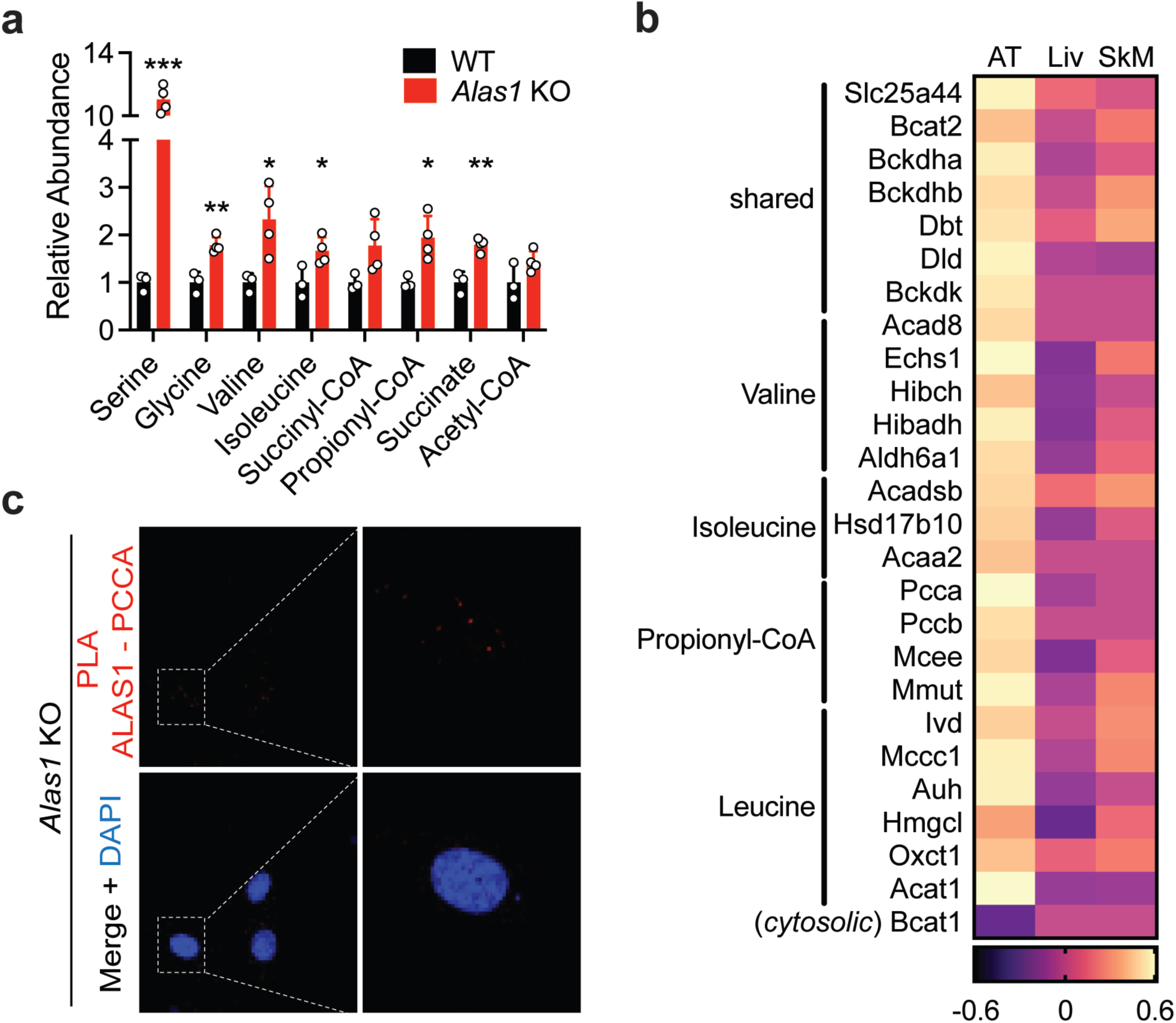
Heme biosynthesis is functionally linked to BCAA and propionyl-CoA metabolism in adipose tissue. **a**) ALAS1 substrates and their precursors accumulate in *Alas1* KO brown adipocytes. **b**) Heatmap showing the correlation in adipose (AT), liver (Liv), and skeletal muscle (SkM) between the expression of *Alas1* and the expression of genes associated with BCAA catabolism. **c**) Proximity ligation assay of ALAS1 and PCCA in *Alas1* KO brown adipocytes. Nuclei counterstain is shown in blue. Data are shown as mean ± SD. * p<0.05, ** p<0.01, *** p<0.001 vs WT; multiple two-tailed Student’s *t*-test.

**Extended Data Figure 8.**
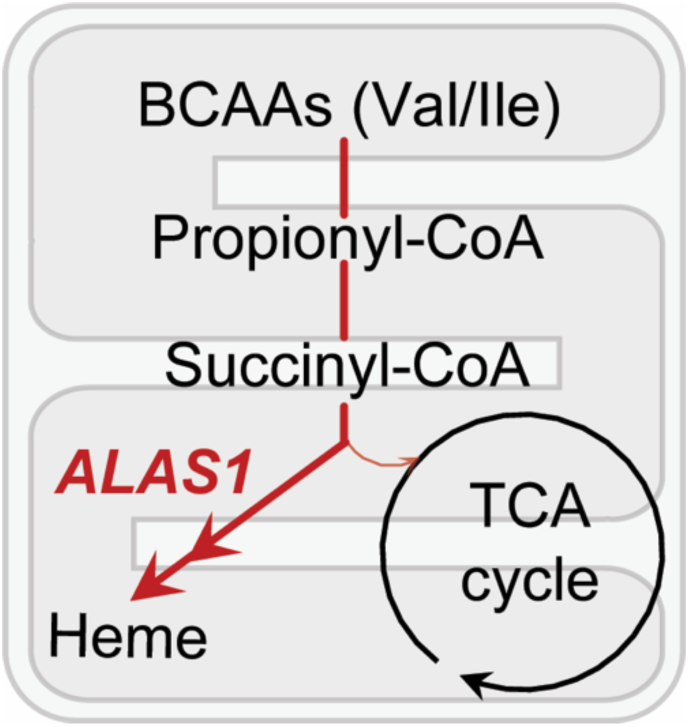
Proposed model whereby brown adipocytes channel BCAA-derived spropionyl-CoA into the heme biosynthetic pathway.

